# DEELIG: A Deep Learning-based approach to predict protein-ligand binding affinity

**DOI:** 10.1101/2020.09.28.316224

**Authors:** Asad Ahmed, Bhavika Mam, Ramanathan Sowdhamini

**Author notes:** **Corresponding author: Email:** (RS). **Co-first author** (Asad Ahmed, Bhavika Mam).

## Abstract

Protein-ligand binding prediction has extensive biological significance. Binding affinity helps in understanding the degree of protein-ligand interactions and has wide protein applications. Protein-ligand docking using virtual screening and molecular dynamic simulations are required to predict the binding affinity of a ligand to its cognate receptor. In order to perform such analyses, it requires intense computational power and it becomes impossible to cover the entire chemical space of small molecules. Recent developments using deep learning has enabled us to make sense of massive amounts of complex datasets where the ability of the model to “learn” intrinsic patterns in a complex plane of data is the strength of the approach. Here, we have incorporated Convolutional Neural Networks to find spatial relationships amongst data to help us predict affinity of binding of proteins in whole superfamilies towards a diverse set of ligands without the need of a docked pose or complex as input. The models were trained and validated using a detailed methodology for feature extraction. We have also tested DEELIG on protein complexes relevant to the current public health scenario. Our approach to network construction and training on protein-ligand dataset prepared in-house has yielded novel insights.

## Introduction

Proteins are a diverse class of dynamic macromolecular structures in living organisms and are essential for the biochemistry and physiology of the organism. Depending on their functional role (s), proteins may bind to other proteins, peptides, nucleic acids and non-peptide ligands with varying affinities. Determining protein-ligand affinity helps in understanding the reaction mechanism and kinetics of the reaction, especially when experimental approaches may not be feasible, and has applications in drug development and pharmacology [1].

Protein-ligand interaction is measured in terms of Binding affinity. The stronger the readout for binding affinity, the stronger the interaction between protein and ligand may be inferred. It is quantified in terms of Inhibition constant (Ki), dissociation constant (Kd), changes in free energy measures (delta G, delta H and IC50) [2]. Predicting binding affinity between a protein and ligand complements experimental approaches and is usually used as a start-point for the latter. Classical prediction methods to score free binding energies of small ligands to biological macromolecules such as MM/GBSA and MM/PBSA typically rely on molecular dynamic simulations for calculations and aid in-silico docking and virtual screening as well as experimental approaches. However, there is a trade-off between computational resources and accuracy [3].

With a recent shift towards the use of machine learning and deep-learning based methods in the field of structural biology, making biologically significant predictions using regression and ‘learning’ intrinsic patterns in a complex plane of available data has led to resource-optimal predictions without compromising on accuracy. Deep learning has been known to learn representations and patterns in complex data forms. Our aim was to apply deep learning to predict binding affinity of protein-non-peptide ligand interaction without the need of a docked pose as input.

Convolutional Neural Networks (CNN) are deep neural networks that use an input layer, output later as well as convolutional hidden layer(s). The first CNN was incorporated by LeCunn in 1998 [4] the connectivity pattern of which was inspired by the elegant experiments of Hubert and Weisel on the mammalian visual cortex in the 1960s [5]. With the growing technical advancements and massive amounts of data, CNNs have emerged popular in biological fields in the recent decade with various applications [6].

In our study, we have used CNNs to provide a quantitative estimate of protein-ligand binding using various sets of features corresponding to protein and ligand respectively by finding spatial relationships amongst the data **without using docked poses as input**. Our approach was validated using ligand-bound complexes from kinases superfamily in the PDB. Kinases belong to a class of enzymes required for substrate-dependent phosphorylation. They are represented across diverse cellular functions like signaling, differentiation, glycolysis [7]. We have also tested our model on COVID-19 main protease [8] of the novel coronavirus strain complexed with various inhibitors of which binding affinities have not been predicted or experimentally determined so far.

## Materials and Method

### Novel Dataset: Raw Data

The raw data for our novel database was obtained from RCSB PDB (9) database, where following were selected as the query parameters.

- **Chain Type:** Protein Chain, No DNA or RNA or DNA/RNA Hybrid.
- **Binding Affinity:** Kd or Ki value present.
- **Chemical Components:** Has ligand (s)
- **X-ray crystallography method:** Resolution upto 2.5 A.

These criteria resulted in a list of 5464 protein PDB IDs, 2568 complexed ligand (s) and corresponding binding affinity values. The search results include the structures present in PDBdatabase, PDBBind (10, 11, 12), PDBMoad (13, 14) and scPDB (15) for its results. Initial raw data database created contained protein structures in PDB format, protein sequences in FASTA format, ligand in SDF format and binding affinity values of corresponding protein-ligand pairs for *5464* complexes.

### Dataset Refinement

The PDB, FASTA and SDF files filtered were further processed to refine our novel dataset, as shown in Figure 1. Protein-ligand complexes were 5,464 in number and corresponded to 29,650 complex **unique** chain-ligand pairs. Binding affinity values were obtained from the RCSB database and protein chain-ligand pairs with corresponding binding affinity as 0 were discarded to reduce statistical errors. This narrowed down the total complexes to 4,750 protein-ligand pairs.

**Figure 1:**
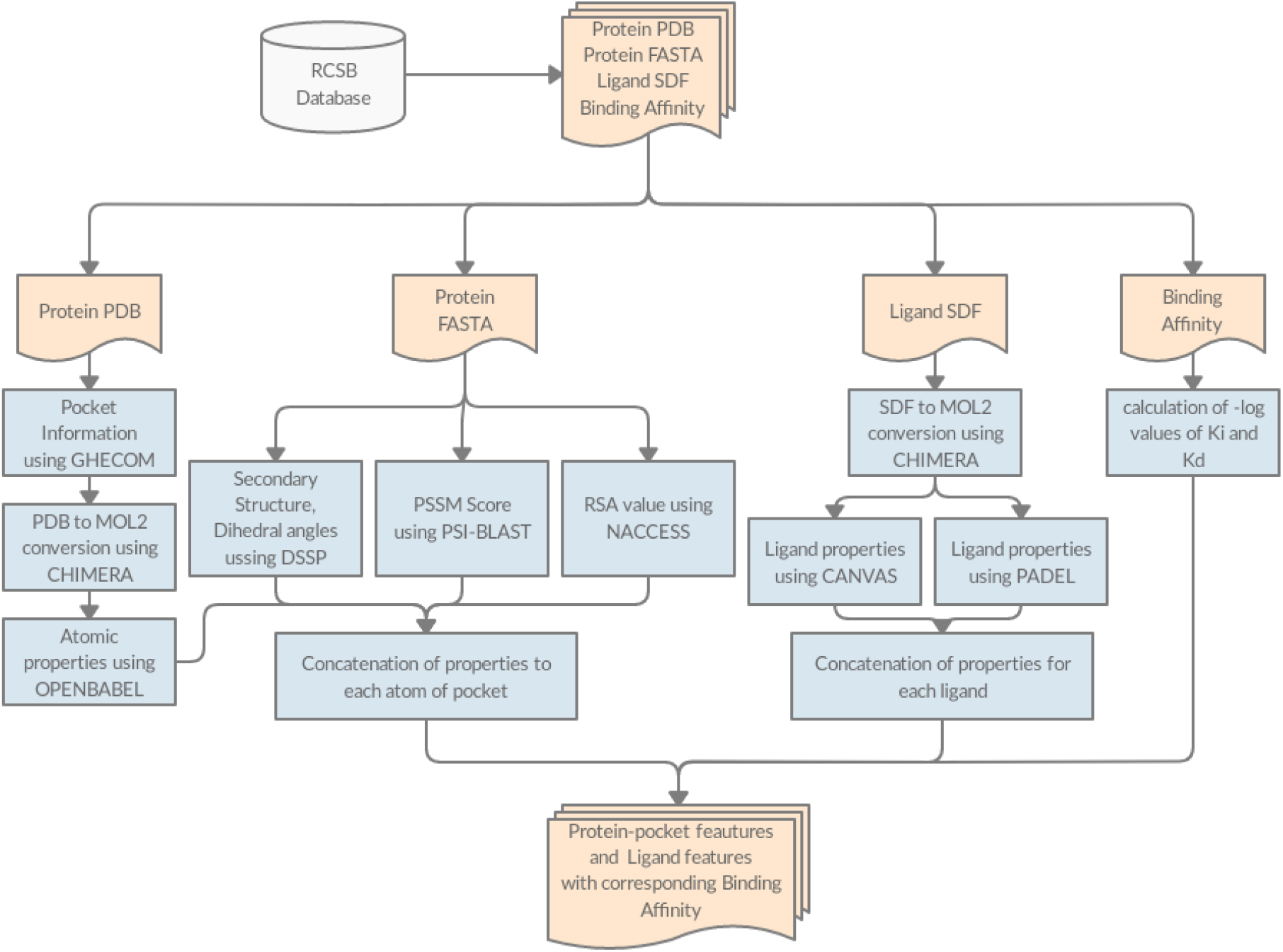
Feature Extraction pipeline.

Pocket information was extracted from the protein using Ghecom (16) and converted to MOL2 format using Chimera (17), which narrowed our results to 4699 pocket-ligand pairs. It narrowed down the size of the dataset to 4286 pocket-ligand pairs. We discarded other protein-ligand pairs with missing PSSM profiles, secondary structure or dihedral angle information.

It resulted in a total of 4041 pocket-ligand pairs, which corresponds to 7414 pocket-ligand pairs containing unique chains.

### Feature Extraction

Training the deep learning network on raw information is known to result in longer time for convergence and less accuracy. We followed a conventional methodology for feature extraction and used the deep learning framework to learn the interaction between the protein-pocket and ligand for their affinity prediction.

### Protein-Pocket features

A comprehensive two-level feature extraction methodology, one at the atomic level and the other at the level of amino acids utilizing structural information and protein sequence respectively.

#### Atomic Level (19 Bits)

- 9 Bit 1 hot or all null hot encoding for atom types: B, C, N, O, P, S, Se, halogen and metal.
- 1 integer for hybridization
- 1 integer representing the number of bonds with heavy atoms
- 1 integer representing the number of bonds with hetero atoms
- 5 bits (1 if present) encoding properties defined with SMARTS patterns: hydrophobic, aromatic, acceptor, donor and ring
- 1 float for partial charges
- 1 integer to distinguish between ligand as −1 and protein as 1

#### Amino Acid level (25 Bits)

We utilized the sequence information of protein to get more features about the protein pocket-ligand interaction.

- Position-Specific Scoring Matrix (PSSM): PSSM is a matrix that represents the probability of mutation at each point of the sequence. It gives a 20 bit-probability for each amino acid at each location. PSSM profiles were obtained using PSI-BLAST (18) with SwissProt as subject database and E-value threshold as 0.001. Chains with less than 50 amino acids were removed from the input dataset.
- Relative Solvent Accessibility (RSA): It is encoded by 1 bit of information for each amino acid that provides whether it is buried or exposed to the solvent. We set a threshold of 25% in RSA values. RSA was obtained using NACCESS (19).
- Secondary Structure: It is encoded by 1 bit of information about the structure as coil, helix or plate and was predicted using the DSSP (20, 21).
- Dihedral Angles: It is encoded by 2 bits of information with phi / psi angles of each of the amino acids and was predicted using DSSP (20, 21) for obtaining dihedral angles.

### Ligand Features

Standard ligand features were calculated for ligands in our dataset using PADEL (22) and 1D, 2D and chemical fingerprints, which includes hybridisation, atom pair interaction, counts of various functional group.

We also used QikProp (34) and QIKPROP (23) to derive ADMET (Absorption, Distribution, Metabolism, Excretion, and Toxicity) properties, which includes the physical properties, solubility and partition coefficients. The exhaustive list of every property calculated is given in the appendix.

It results in a 1D array of 14,716 dimensions containing the various properties of a given ligand. This is used as a feature vector representing the ligand represented in MOL2 format.

### Grid Formation

The three-dimensional co-ordinates of atoms were converted into a 3D grid of resolution 10Å with 1Å spacing between the two axes centered along the centroid of the ligand. Atoms outside each such grid were discarded. The atoms lying inside the grid were rounded up to the nearest coordinate of the grid where features of corresponding atoms that lay in the same coordinates were added up.

This resulted in projecting ligand-interacting residues into a three-dimensional cube with features representing the atomic as well as protein-based properties of each atom of the protein pocket.

### Strategies

Detailed and complete block diagrams with inputs are provided in Figures 2, 3 as well as in Supplementary Materials.

**Figure 2:**
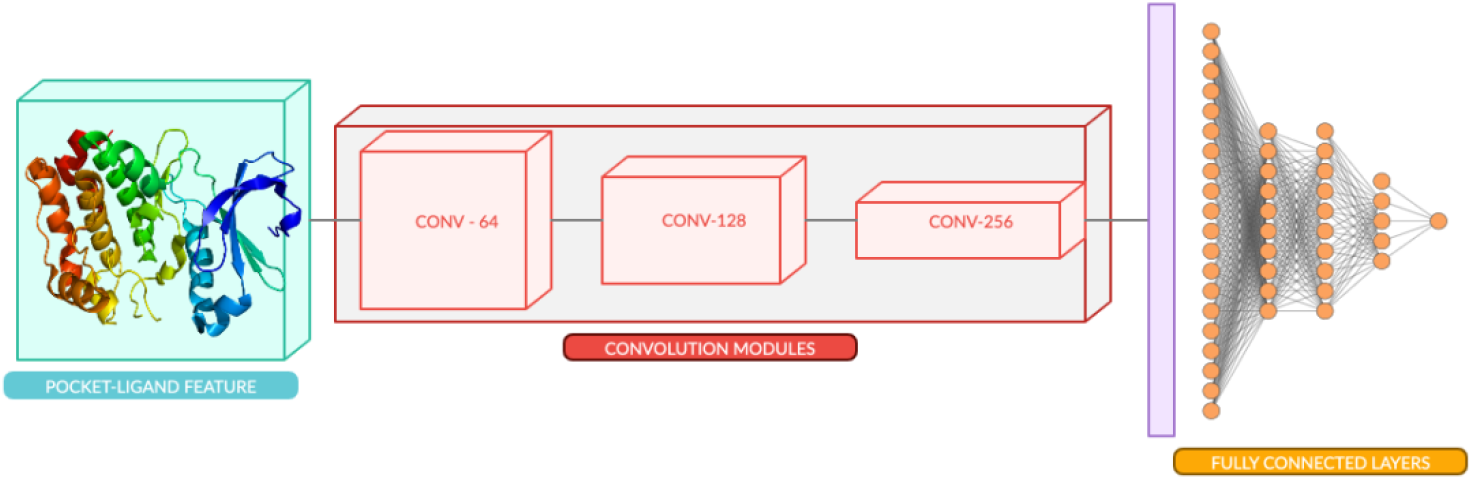
Training framework for Atomic Model. The framework is trained on 19 bits features each of protein-pocket and ligand together as input.

**Figure 3:**
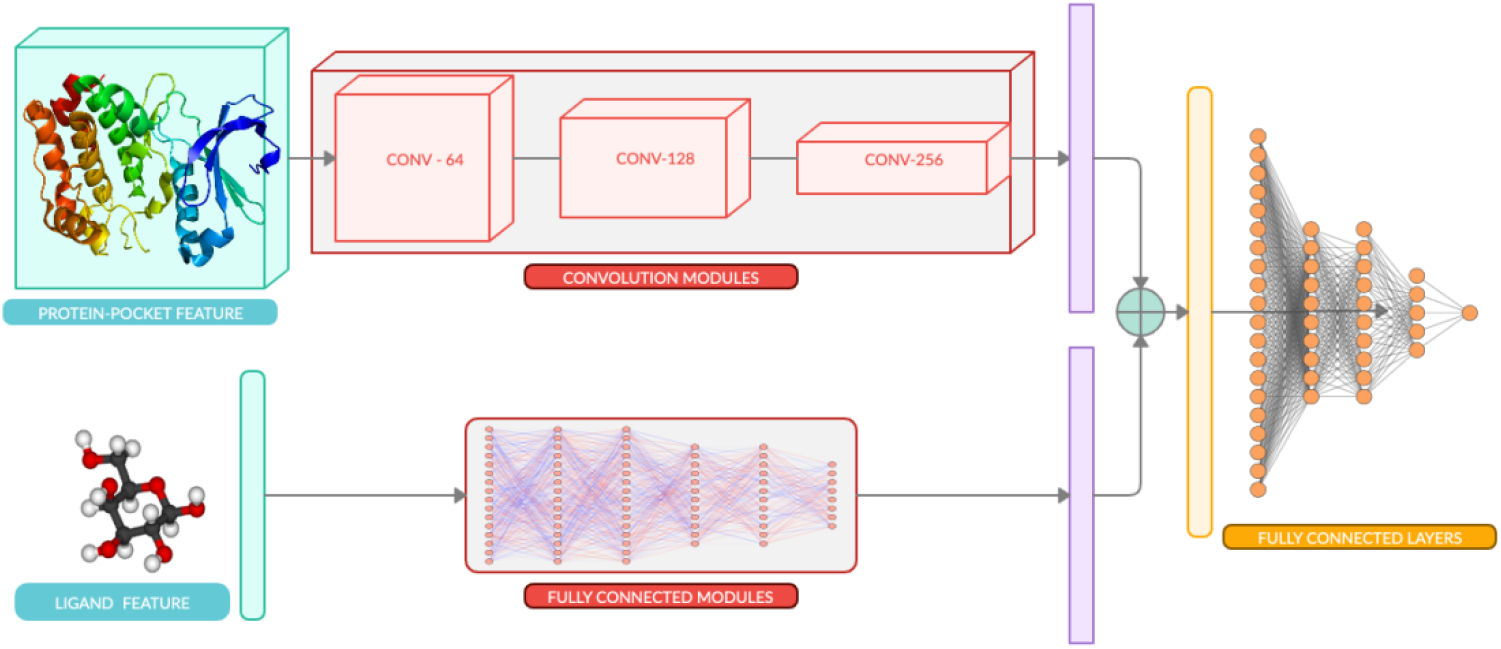
Training framework for Composite Model. The framework is trained on 44 bits features of protein-pocket and 14716 bits of ligand as separate inputs.

### Atomic Model

#### Preprocessing

Features were calculated at the atomic level (Section 4.1.1) corresponding to each atom of an amino acid and ligand. A 19-bit vector was calculated that uniquely identified each of the atoms in the 3D co-ordinates of a given protein-pocket and ligand complex. A 4D tensor each of size m x m x m x 19, i.e. the 3 coordinates (x, y, z) and the features, where m represents the number of atoms present in a complex was constructed as the feature vector representing the given protein pocket-ligand.

The 4D vector contains the protein-pocket features and was converted to a 3D grid using grid featurization (Section 4.3). The 3D- featurized grid is essentially a 4D tensor, where the coordinates are approximated to the points on the grid.

The dataset is converted to vectors and is divided into training:validation:test sets in ratio 80:10:10.

### Architecture

Convolutional Neural Networks (24) have been used to capture spatial features in an image. We use CNNs to capture the interaction between ligand and protein atoms in three-dimensional space. A network was constructed (Figure 2) with a 3D CNN of varying channel sizes of [64, 128, 256] with non-linear activation ReLU after each layer, each 3D CNN had a filter of 5Å cube which was used to perform convolution operations. MaxPool (25) layer that acts in three dimensions to lower the dimension with a pool size of 2Å cube and Batch Normalization (26) layer is added after each CNN layer, this in turn decreases the training time and helps in faster convergence.

The latent features learnt from the above CNN layers were then flattened and used for calculating the binding affinity of the protein pocket-ligand pair. The CNN derives the relation among the 3D coordinates and their features, which would correspond well to the binding affinities of complexes.

The features from the last CNN layer are then flattened out, and passed through a fully connected neural network having the number of neurons as [1000, 500, 250] with ReLU as non-linearity after each layer. Dropout (25) is added after each layer to prevent overfitting by forcing the neural network to learn various other pathways by randomly assigning neurons to zero, 0.50 as Dropout threshold. Dense network predicts a regressive value of Binding Affinity, corresponding to a single neuron output. Training framework is shown in **Figure 2** and a detailed layer network is shown in **Figure 4 (a)**.

### Training

The featurized protein-pocket grid formed was rotated to all 24 combinations possible, such that the network is able to learn in an orientation invariant form.

The network was trained by taking Mean Square error between the predicted and actual values as a loss function. The network was optimized using Adam (27) as the optimizer with a learning rate of 1e-5 and weight decay of 0.001 for 20 epochs. Network was trained on an Nvidia Pascal GPU using Pytorch (28) as the framework.

### Composite Model

#### Preprocessing

Features were calculated at the amino acid level (Section 4.1.2) and were concatenated alongside the atomic level features (Section 4.1.1) to each atom of amino acid. It results in a 44-bit vector uniquely identifying each of the atoms in the 3D co-ordinates of a given protein. A 4D tensor each of sizes m x m x m x 44, i.e. the 3 coordinates (x, y, z) and the features, where m represents the number of atoms present in a complex is constructed as the feature vector of protein pocket.

The 4D vector contains the protein-pocket features, it was converted to a 3D grid using grid featurization (Section 4.3). The 3D featurized grid is essentially a 4D tensor, where the coordinates are approximated to the points on the grid.

The ligands were separately featurized by calculating the ligand properties (Section 4.2), which results in a 1D tensor.

The dataset is converted to vectors and is divided into training:validation:test sets in ratio 80:10:10.

### Architecture

A multi-input network was constructed (29) with a 3D CNN (24) of varying channel sizes of [64, 128, 256] with non-linear activation ReLU after each layer, each 3D CNN had a filter of 5Å cube which was used to perform convolution operations. We also added MaxPool (25) layer that acts *three-dimensionally* to lower *dimensionality* while retraining features learnt after each CNN layer. It has a filter size of 2Å cube. Batch Normalization (26) layer was added after each CNN module for faster convergence.

The ligand features were passed through the dense layers of sizes [7000, 5000, 2000] with ReLU as non-linearity after each layer and we also perform dropout operations after each dense layer to prevent it from overfitting (30). This results in a latent vector representing the relevant features for each ligand.

The latent output from the CNN layers is flattened and concatenated with the latent feature vector of ligand, to create one single feature vector of protein pocket-ligand interactions. This vector is passed through a densely connected neural network having the number of neurons as [7000, 2000, 500, 200] with ReLU as non-linearity after each layer and we used Dropout after each layer also to prevent overfitting forcing the neural network to learn various other pathways by randomly assigning weights of neurons to zero, with 0.50 as Dropout threshold. This dense network finally predicts a regressive value of Binding Affinity, corresponding to a single neuron output.

Training framework is shown in Figure 3 and a detailed layer network is shown in Figure 4 (b)

### Additional Case studies of specific protein families

Recently deposited complexes of COVID-19 main protease with various inhibitors deposited in the PDB were used for the purpose of our study (**Table 3**). The crystal structure complexes (PDB IDs: 5R7Y, 5R7Z, 5R82, 5R84) of the COVID-19 main protease with inhibitors ((Z45617795: N-[(5-methylisoxazol-3-yl)carbonyl]alanyl-L-valyl-N~1~-((1R,2Z)-4-(benzyloxy)-4-oxo-1-{[(3R)-2-oxopyrrolidin-3-yl]methyl)but-2-enyl)-L-leucinamide); Z1220452176: (~{N)-[2-(5-fluoranyl-1~{H)-indol-3-yl)ethyl]ethanamide); Z219104216: 6-(ethylamino)pyridine-3-carbonitrile; Z31792168:2-cyclohexyl-~{N)-pyridin-3-yl-ethanamide)) respectively has been recently deposited in PDB (2020; unpublished).

Another study has deposited the complex of the COVID-19 main protease with a broad-spectrum inhibitor X77 (N-(4-tert-butylphenyl)-N-[(1R)-2-(cyclohexylamino)-2-oxo-1-(pyridin-3-yl) ethyl]-1H-imidazole-4-carboxamide) (2020; unpublished).

In order to compare affinity of deoxycholate with homologous proteins of the periplasmic C-type cytochrome (**Table 4**), Ppc homologs PpcA (PDB: 1OS6), PpcB (PDB: 3BXU), PpcC (PDB: 3H33), PpcD (PDB: 3H4N) and PpcE (PDB: 3H34) and ligand deoxycholic acid (Pubchem CID: 222528) were gathered. These were processed and DEELIG was used to predict the binding affinity of each homolog with the ligand.

### Training

The featurized protein-pocket grid formed was rotated to all 24 combinations possible, such that the network is able to learn in an orientation invariant form.

The featurized protein pocket-ligand pair of training set was passed through corresponding the network and trained by taking Mean Square error between the predicted and actual values as a loss function. The network was optimized using Adam (27) as the optimizer with a learning rate of 1e-5 and weight decay of 0.001. The network was trained on an Nvidia Pascal GPU using Pytorch (28) as the framework.

### Performance Evaluation

The predicted value of our regression-based approach is the negative natural logarithmic value of Kd or Ki. This is then converted to its antilog to obtain Kd or Ki value in nanoMolar quantity.

The performance of the models was quantified using Mean Absolute error (MAE) and Root mean square error (RMSE). It was tested on validation and testing sets which were initially divided from our dataset as mentioned in the training section. Lower error corresponds to better learning capacity of the model. Standard deviation among the real and predicted values was also calculated.

The MAE, RMSE and SD values are shown in Table 1.

**Table 1:** Predictions accuracy on test set of our novel dataset.

| Method | MAE | RMSE | SD |
| --- | --- | --- | --- |
| Atomic Model | 2.84 | 3.93 | 2.62 |
| **Composite Model** | **2.27** | **3.07** | **2.06** |

For the purpose of training and testing models, one NVIDIA Tesla P100 GPU cluster was used. Computational time taken for featurization of the dataset, training and testing were 52 hours, 22 hours and 8 minutes respectively.

## Results and Discussion

Two modules were trained. The first module was trained using a small set of features for protein and ligand, which were represented together in a 3D grid space. This approach has also been part of a previous study (29). However, the previous study uses a restricted ligand set that does not involve larger ligands. Here we have used a diverse set of ligands as one of our inputs. With training of Atomic Model for *35 epochs*, ***MAE score of 2.84 was achieved*** (Table 1).

We constructed another module that enabled us to improve on the ligand and protein based information. To this purpose, we used an increased feature vector size which amounted to 14716 bits in size for ligand and 44 bits for each atom of protein. With training of Composite Model for only *4 epochs*, **MAE score of 2.27** was achieved (Table 1).

The performance of our model was further evaluated using ligand-bound complexes from the kinase superfamily from PDB. The composite model outperformed the atomic model significantly and with lower standard deviation. (**Table 2**).

**Table 2:** Predictions accuracy on kinases.

| Method | MAE | RMSE | SD |
| --- | --- | --- | --- |
| Atomic Model | 2.48 | 3.24 | 3.11 |
| **Composite Model** | **2.24** | **2.71** | **2.67** |

In light of the ongoing coronavirus pandemic, we tested protein-ligand complexes from the coronavirus (CoV) family. The COVID-19 main protease is a key enzyme for the novel strain of coronavirus that is being implicated in the pandemic. A recent study involved testing of in-vitro binding efficacy of coronavirus COVID-19 virus main protease (Mpro) with a potent reversible synthetic inhibitor, N3 (31). However, the highly potent inhibition by N3 rendered the experimental determination of binding affinity not achievable. Using the structure of Mpro at high resolution (7BQY: 1.7 Angstrom), we have been able to predict the binding affinity of N3 to 3.1e+4 nanomolar (Table 3). This value agrees with the observed high affinity in the course of recent experiments (31).

**Table 3:** Predictions of Binding Affinity on COVID-19 complexes.

| PDB | Ligand | -Log (Kd/Ki) | [Kd] or [Ki] (nM) |
| --- | --- | --- | --- |
| 5R7Y | Z45617795 | 11.96 | 6.39e+3 |
| 5R7Z | Z1220452176 | 7.69 | 4.57e+5 |
| 5R82 | Z219104216 | 6.12 | 2.18e+6 |
| 5R84 | Z31792168 | 8.32 | 2.43e+5 |
| 6W63 | X77 | 15.34 | 2.17e+2 |
| 7BQY | N3 | 10.38 | 3.10e+4 |

We used complexes of COVID-19 main protease with various inhibitors (**Materials and Methods**; **Table 3**) to predict their respective binding affinities as their experimental values have not been made available. Based on our model-based predictions, broad spectrum inhibitor X77 scores for highest affinity followed by ligands Z45617795, N3, Z31792168, Z1220452176 and Z219104216 in the order of decreasing binding affinity (**Table 3**) strengthening the suitability of X77 as a potential candidate against COVID-19 virus protease

A triheme cytochrome from the sulfur-, metal- and radionuclide-reducing bacteria, *Geobacter sulfurreducens,* named PpcA binds strongly to deoxycholate [10]. However, its triheme paralogous counterparts PpcB, PpcC, PpcD and PpcE do not bind to deoxycholate [11, 12]. Our results also predict that ligand deoxycholate binds with high affinity to periplasmic C-type cytochrome A (PpcA) but not to its homologs PpcB, PpcC, PpcD and PpcE (**Table 4**).

**Table 4:** Predictions of Binding Affinity on homologs of Periplasmic C-type Cytochrome (Ppc) family.

| Homolog | PDB ID | Prediction<br>Kd or Ki (uM) |
| --- | --- | --- |
| PpcA | 1OS6 | 4.512 |
| PpcB | 3BXU | 416.042 |
| PpcC | 3H33 | 835.232 |
| PpcD | 3H4N | 483.678 |
| PpcE | 3H34 | 187.157 |

## Conclusion

Deep-learning based approaches have been implemented for prediction of binding affinity. One of the studies used atomic level features of complex in a CNN based framework for binding affinity prediction (35), while another study used protein sequence level features in a CNN based framework for prediction (36). Another approach used as been to use feature learning along with gradient boosting algorithms to predict binding affinity (36). Here, we provide a composite model that incorporates tripartite structural, sequence and atomic level features with those of the atomic and other chemical features of the ligand to predict binding affinity of a putative complex.

We propose a deep-learning based approach to predict ligand (eg., drug)–target binding affinity using only structures of target protein (PDB format) and ligand (SDF format) as inputs. Convolutional Neural Networks (CNN) were used to learn representations from the features extracted from these inputs and hidden layers in the affinity prediction task. We used two approaches to feature extraction-atomic level as well as composite level and compared their performance using the same network. We have trained on complexes from PDB across all taxa filtered as per few starting criteria including crystal quality. Our results are validated and reflected in the performance scores. The baseline to the results of our approach is the study by Stepniewska-Dziubinska et al 2018 [27], the performance of which our study has exceeded (**Results**).

Our algorithm relies on certain inputs including sensitive binding cavity detection by the Ghecom algorithm (Kawabata, 2010) that uses mathematical morphology to find both deep and shallow pockets (if any) in a given protein. The coordinates of the predicted binding cavity of the protein (grid) are rotated to various combinations and are placed around the centroid of the ligand and the resultant 4-D tensor is processed further for features along the CNN (**Materials and Method**). Hence, ligand-bound poses are not used as input. Our dataset has ~5k+ complexes and also includes complexes that were not part of PDBBind (which is usually used to benchmark and is derived from PDB). The ligand set we have used also represents a diverse set (**Supplementary Materials SM Files 1 and 2**) and is one of the highlights of our approach. The predictions from DEELIG can in fact help existing databases like RSCB PDB, PDBMoad and PDBBind in filling missing binding affinity data for complexes.

We have constructed a novel dataset that represents a diverse set of ligands and using a novel deep learning based approach we have achieved significant improvement in prediction of binding affinity of protein-ligand complexes. Interestingly, our approach performed better without ligand coordinates as input. To counter filtering or noise reduction in our dataset, our dataset constructed is smaller than PDBBind (35) but we have overcome the constraints on ligand selection part of a previous study (29). Although our dataset contains 5464 complexes compared to 16,151 complexes found in PDBBind, the ligands used as part of our training include 452 unique ligands absent in PDBBind. This helps in achieving ligand diversity during training the CNN model. The similarity matrix constructed from the binary fingerprints of ligands used in the dataset supports our claim of improved ligand diversity in our dataset (Supplementary File S1).

We have highlighted a few examples such as complexes of kinases and viral drug targets only to reinforce the broader applicability of our approach (**Tables 2 and 3**). Our predictions are in line with experimental observations [32, 33, 34] that deoxycholate binds to PpcA cytochrome but not to homologs PpcB - E cytochrome (**Table 4)**.

We have also eliminated the need of providing ligands in a complex form with protein. Thus a given protein pocket may be tested for the degree of binding for any given ligand. This can be extended to predicting potential binding partners for proteins in other superfamilies as well. It is also important to consider that docking score and pose is not a reliable correlation with MM/GBSA poses (37). **DEELIG can be used for a member of any protein superfamily and a non-peptide ligand, the docking pose of which may or may not be known.**

The code repository for the project is publicly available at: https://github.com/asadahmedtech/DEELIG

## Future Direction

Binding affinity predictions through DEELIG can be extended to protein-ligand complexes of protein superfamilies where the affinity is quantitatively unknown due to experimental limitations or where the potential for binding is yet to be explored *in vitro*. A webserver to implement DEELIG for easy online access would be useful for the general scientific community and this will also be in the pipeline. A later version of DEELIG which is trained on peptide ligand dataset will also be worked on.

## Supporting information

Compressed folder with raw data of ligands used in analysis.

## Supplementary Files

a. **SM compressed folder:** dataset.gz (can be retrieved from https://drive.google.com/file/d/1JE3gQuTXprRVghygAwR9HESvABHKED0L/view?usp=sharing)
b. **SM Files for Ligand diversity analysis:** Similarity matrix (**SM File 1**) and clustering (**SM File 2**) of unique ligands-https://drive.google.com/drive/folders/1Ar64qn8vD0sSdPWptPgOkPeM7pWghKi7?usp=sharing
c. **SM File 3:** Dataset_distribution.xls
d. **SM File 4:** Dataset_details.xls

## Appendix

## A.1 Property list for ligand features

Following properties of ligand were calculated using PADEL (22),

- Basic Group Count
- Carbon Type
- Hybridization Ratio
- Manhold LogP (The Ratio of carbon to hetero atoms)
- Number of Aromatic bonds
- MACCSS Key
- Klehotaroth fingerprints (Types and Counts)
- AtomPair2D fingerprints (Types and Counts)

Following are ADMET and present in PADEL

- *donorHB*
- *accptHB*
- *Constitutional (Electronegativity)*
- *rotatableBondCounts (#ringatoms)*
- *RuleofFive*
- *VABC (Volume)*
- *Weight (mol_MW)*

Following {*ADMET*) properties of ligand were calculated using QikProp (34) and QIKPROP (16, 21),

- Amine
- Amidine
- Acid
- Amide
- Rotor
- rtvFG (reactive functional groups)
- mol_MW, dipole
- Volume
- donorHB
- accptHB
- QPpolrz (polarizability)
- SASA

- SASA (probe of 1.4A)
- FOSA (hydrophobic component of SASA)
- FISA (hydrophilic component of SASA)
- PISA (pi of SASA)
- WPSA (polar of SASA)
- SAFluorine
- SAamideO
- **Partition coefficients =>**

○ QPlogPC16
○ QPlogPoct
○ QPlogPw
○ QPlogPo/w
- CIQPlogS (**Conformation indie aqueous solubility**)
- IP (ev) (ionization potential)
- EA (eV) (electron affinity)
- #metab (likely metabolic reactions)
- PSA (van der waals SA of polar N and O atoms)
- #NandO, #ringatoms (number of atoms in rings)
- #in34 (number of atoms in 3 or 4 membered rings)
- #in56 (number of atoms in 5 or 6 membered rings)
- #noncon (ring atoms cannot form conjugated aromatic bonds)
- #nonHatm (heavy atoms- nonhydrogen atoms)
- RuleOfThree
- RuleOfFive (lipinski violations)
- QPlogKhsa (binding to human serum albumin)
- PercentHuman-OralAbsorption
- Globular nature index

## A.2 Network Layout for modules

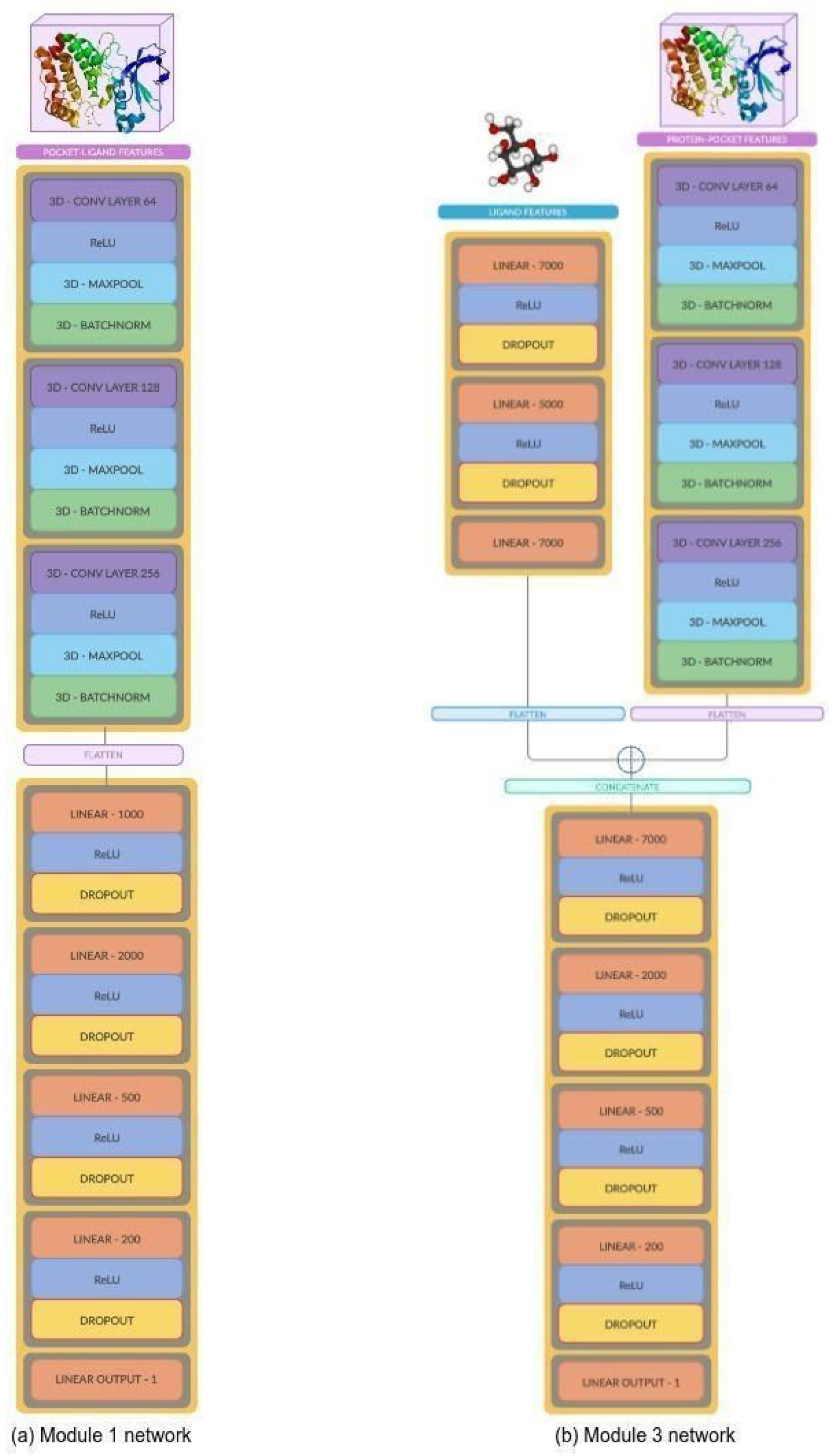

## B.1 Kinases prediction dataset

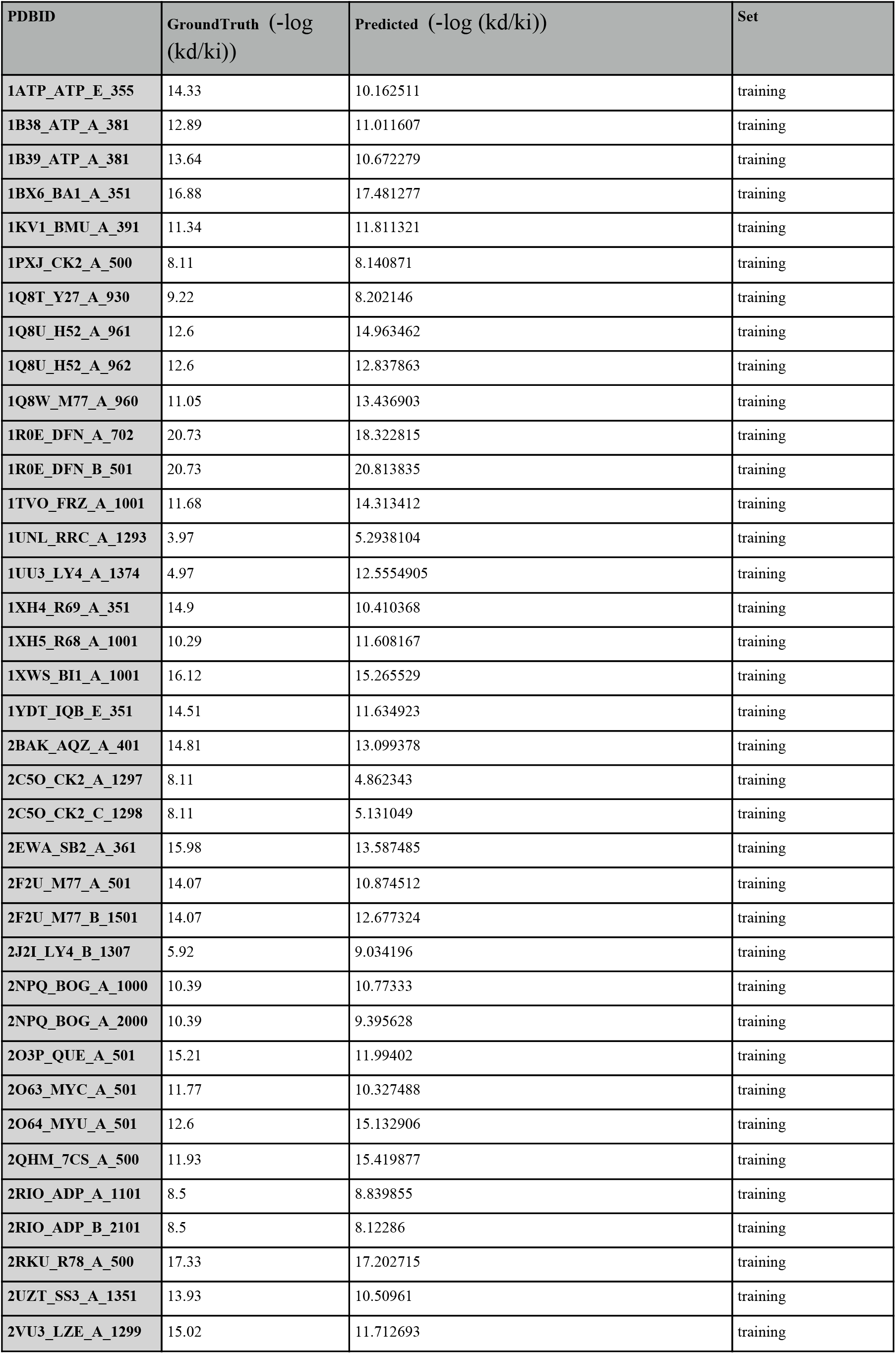
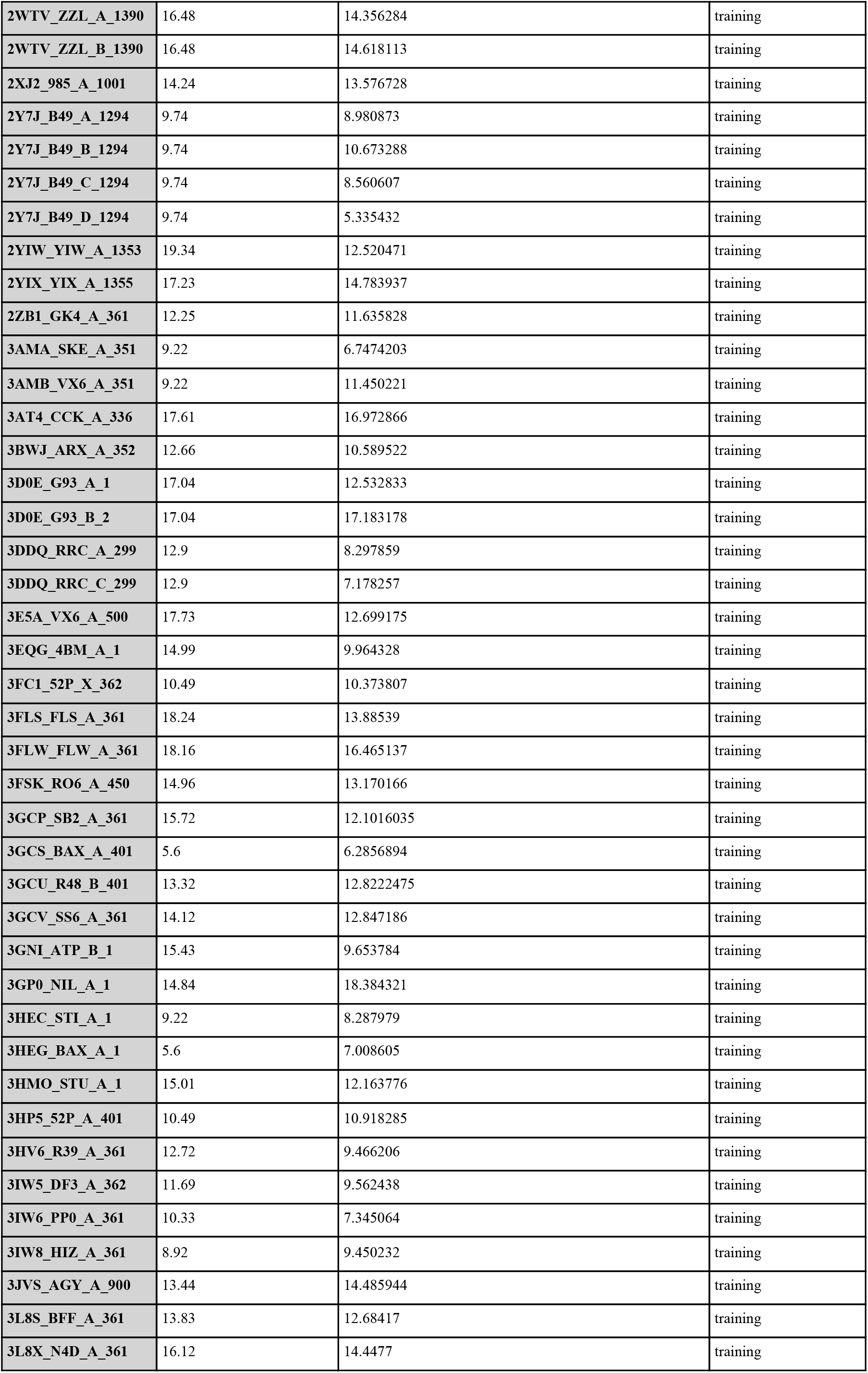
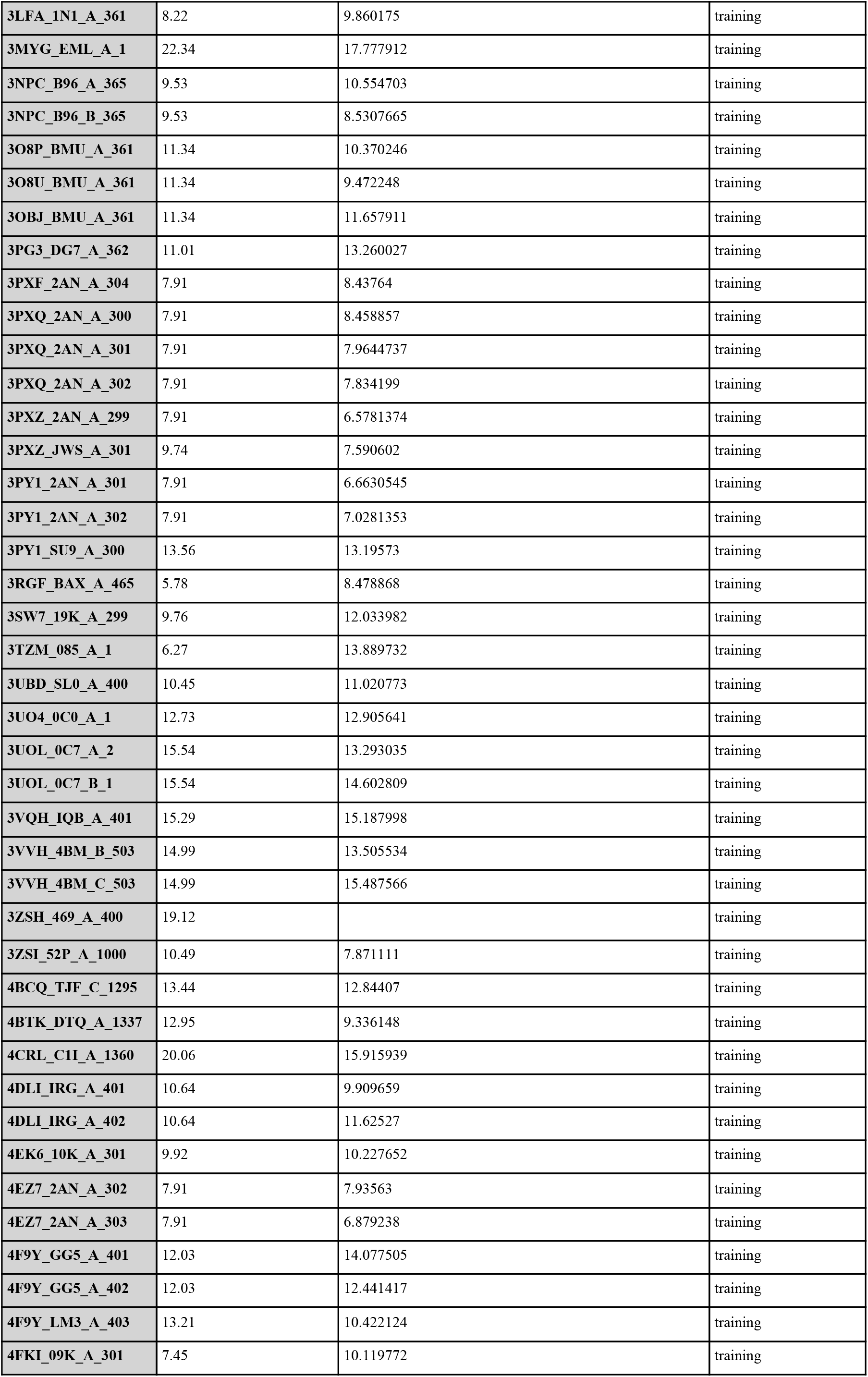
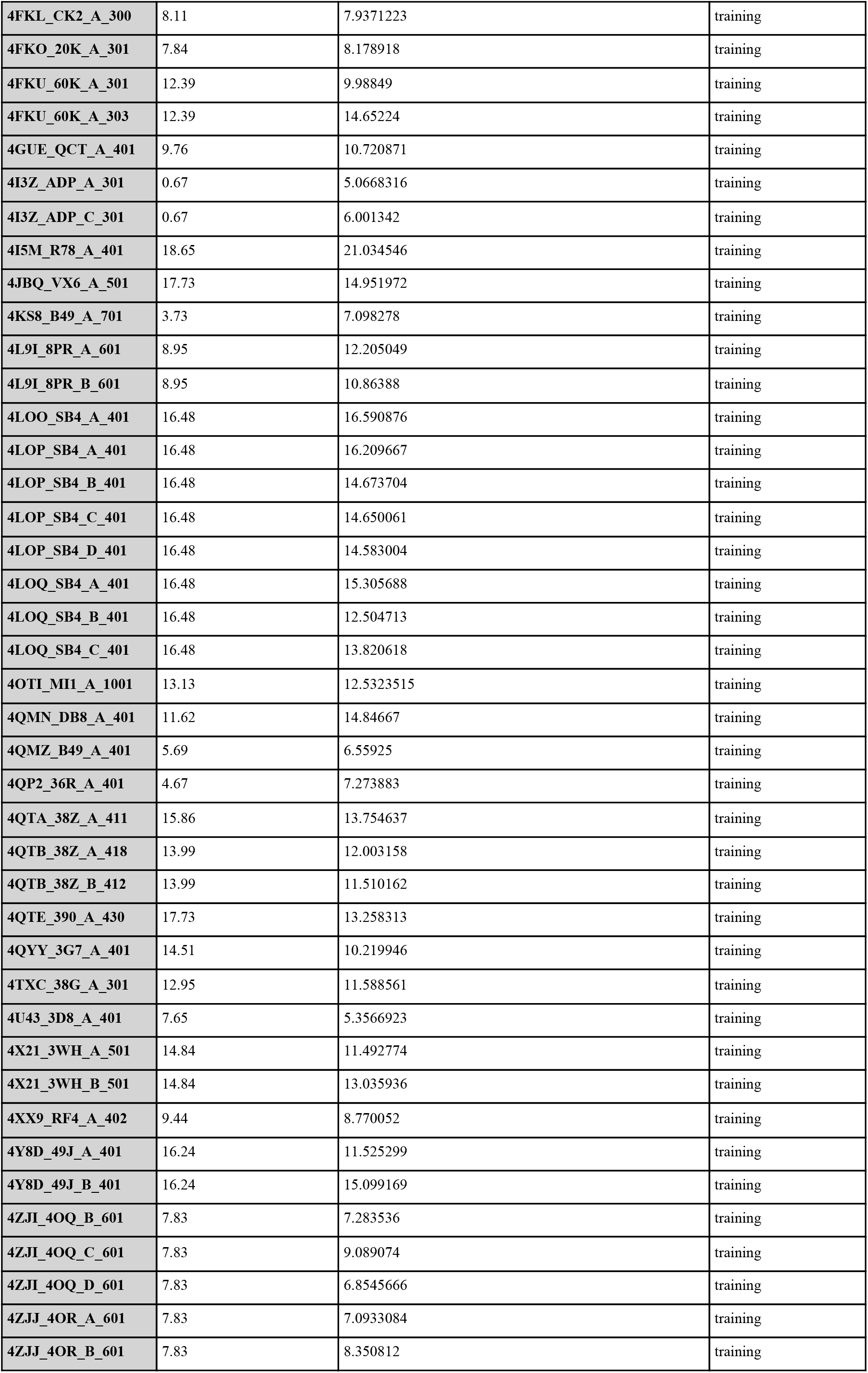
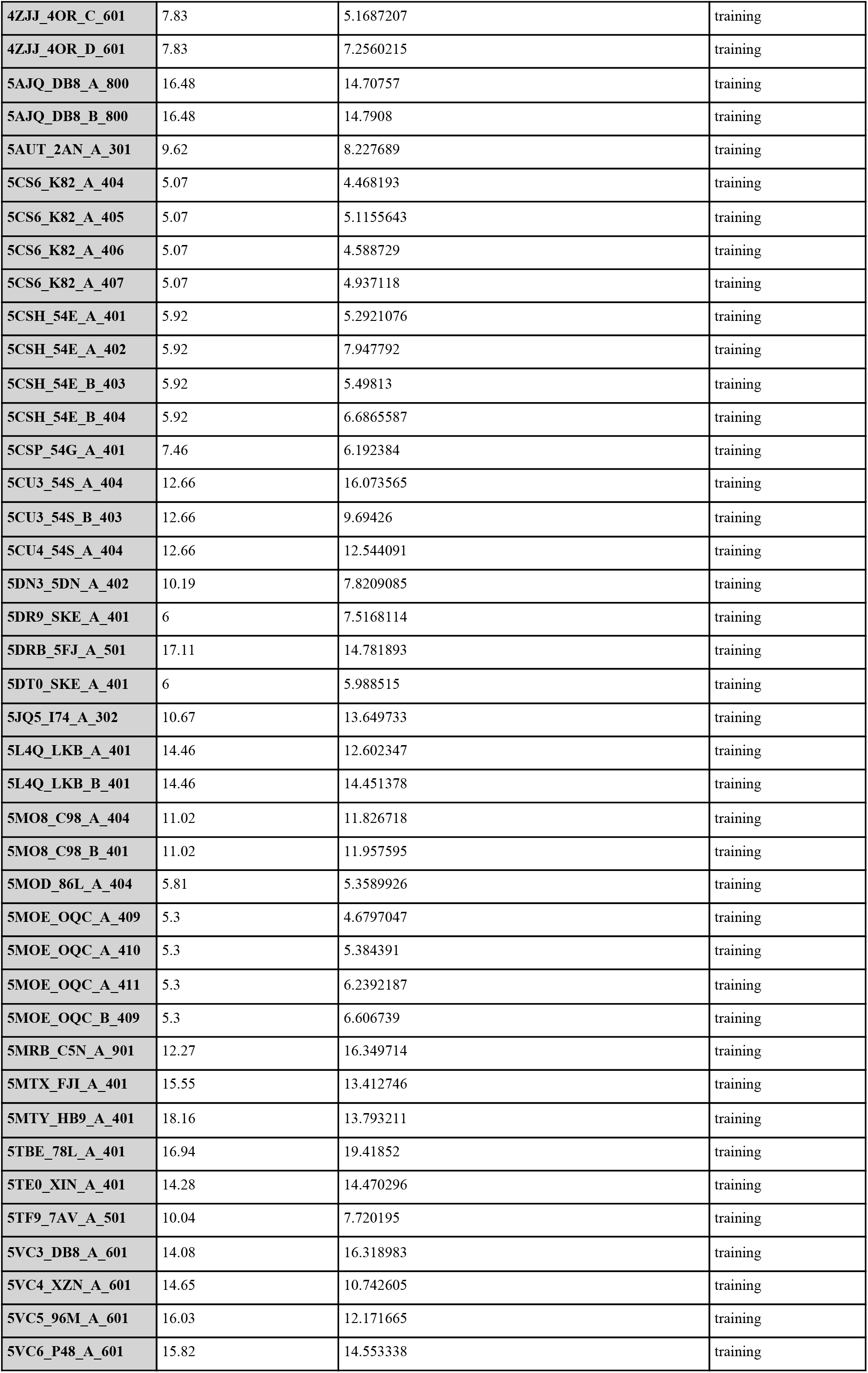
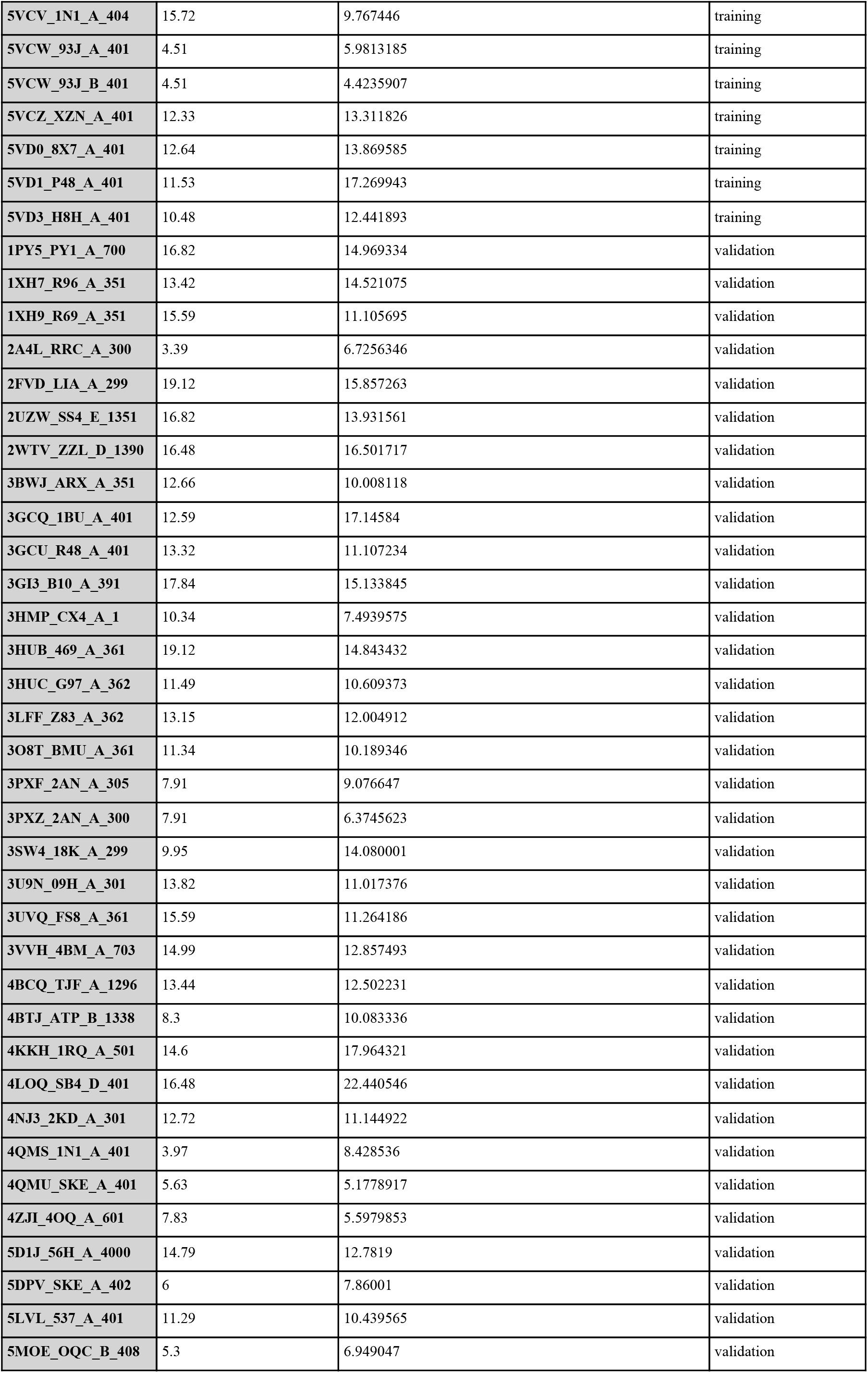
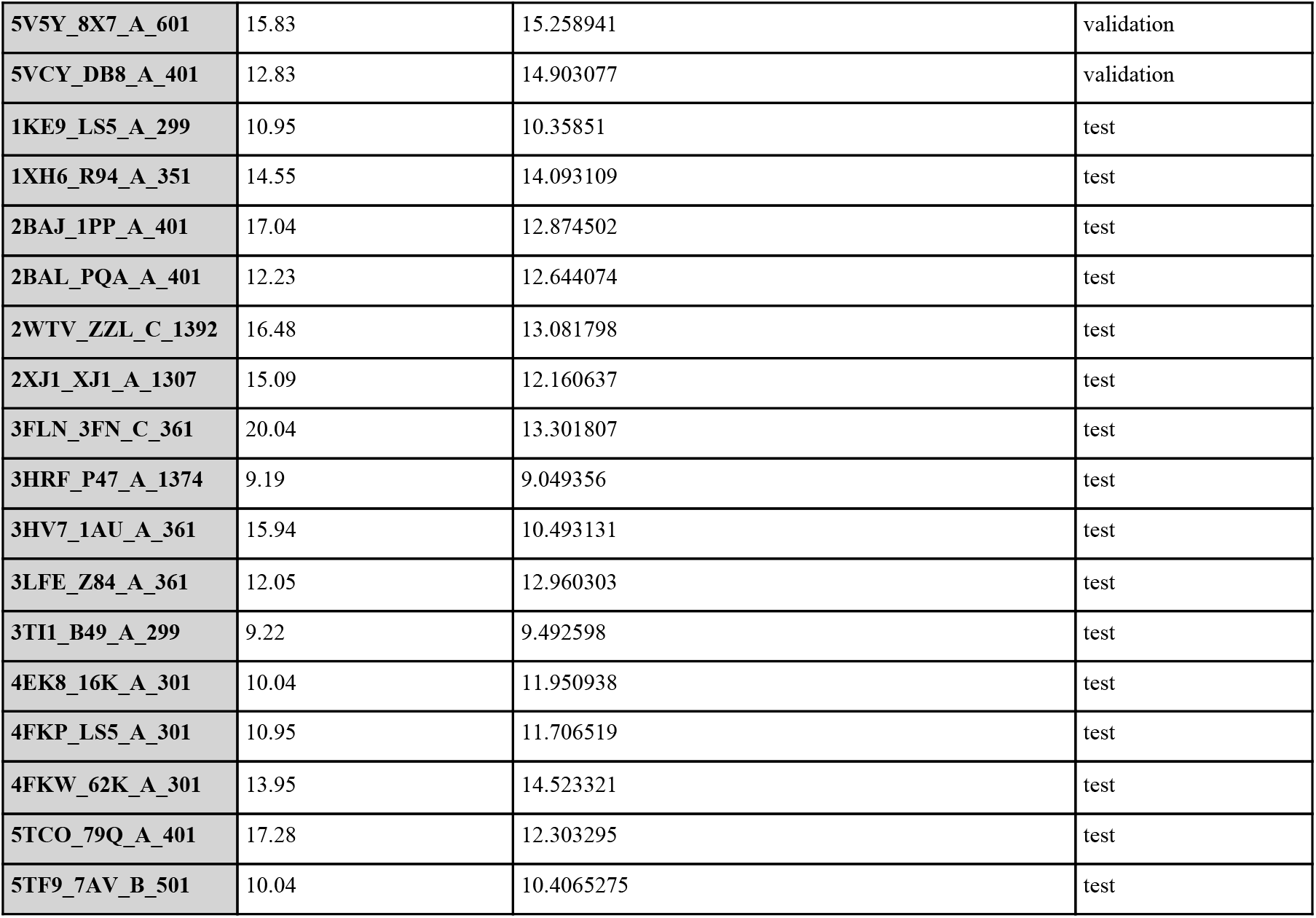

## Acknowledgements

AA acknowledges funding awarded by the Indian Academy of Sciences, Bangalore (2019). BM would like to acknowledge Tata Trusts-TDU Fellowship for PhD awarded to her from 2017 to 2019. All authors acknowledge NCBS for infrastructural support.

## Author Contributions

**Conceptualization:** Bhavika Mam, Asad Ahmed

**Data curation:** Asad Ahmed, Bhavika Mam

**Formal analysis:** Asad Ahmed, Bhavika Mam

**Funding acquisition:** Ramanathan Sowdhamini.

**Investigation:** Asad Ahmed, Bhavika Mam

**Methodology:** Asad Ahmed, Bhavika Mam

**Project administration:** Ramanathan Sowdhamini

**Resources:** Ramanathan Sowdhamini

**Supervision:** Ramanathan Sowdhamini

**Validation:** Asad Ahmed, Bhavika Mam

**Visualization:** Asad Ahmed

**Writing –** Asad Ahmed, Bhavika Mam

**Writing – review & editing:** Ramanathan Sowdhamini

